# A connectome-based, corticothalamic model of state- and stimulation-dependent modulation of rhythmic neural activity and connectivity

**DOI:** 10.1101/697045

**Authors:** John D Griffiths, Anthony Randal McIntosh, Jeremie Lefebvre

## Abstract

Rhythmic activity in the brain fluctuates with behaviour and cognitive state, through a combination of coexisting and interacting frequencies. At large spatial scales such as those studied in human M/EEG, measured oscillatory dynamics are believed to arise primarily from a combination of cortical (intracolumnar) and corticothalamic rhythmogenic mechanisms. Whilst considerable progress has been made in characterizing these two types of neural circuit separately, relatively little work has been done that attempts to unify them into a single consistent picture. This is the aim of the present paper. We present and examine a whole-brain, connectome-based neural mass model with detailed long-range cortico-cortical connectivity and strong, recurrent corticothalamic circuitry. This system reproduces a variety of known features of human M/EEG recordings, including a 1/f spectral profile, spectral peaks at canonical frequencies, and functional connectivity structure that is shaped by the underlying anatomical connectivity. Importantly, our model is able to capture state-(e.g. idling/active) dependent fluctuations in oscillatory activity and the coexistence of multiple oscillatory phenomena, as well as frequency-specific modulation of functional connectivity. We find that increasing the level of sensory or neuromodulatory drive to the thalamus triggers a suppression of the dominant low frequency rhythms generated by corticothalamic loops, and subsequent disinhibition of higher frequency endogenous rhythmic behaviour of intra-columnar microcircuits. These combine to yield simultaneous decreases in lower frequency and increases in higher frequency components of the M/EEG power spectrum during states of high sensory or cognitive drive. Building on this, we also explored the effect of pulsatile brain stimulation on ongoing oscillatory activity, and evaluated the impact of coexistent frequencies and state-dependent fluctuations on the response of cortical networks. Our results provide new insight into the role played by cortical and corticothalamic circuits in shaping intrinsic brain rhythms, and suggest new directions for brain stimulation therapies aimed at state-and frequency-specific control of oscillatory brain activity.

**Author Summary:** One of the most distinctive features of brain activity is that it is highly rhythmic. Developing a better understanding of how these rhythms are generated, and how they can be controlled in clinical applications, is a central goal of modern neuroscience. Here we have developed a computational model that succinctly captures several key aspects of the rhythmic brain activity most easily measurable in human subjects. In particular, it provides both a conceptual and a concrete mathematical framework for understanding the well-established experimental observation of antagonism between high- and low-frequency oscillations in human brain recordings. This dynamic has important implications for how we understand the modulation of rhythmic activity in diverse cognitive states relating to arousal, attention, and cognitive processing. As we demonstrate, our model also provides a tool for investigating and improving the use of rhythmic brain stimulation in clinical applications.

## Introduction

A key characteristic of the fluctuations in extracranial electrical and magnetic fields measured by electroencephalography (EEG) and magnetoencephalography (MEG), resulting from the collective activity of large numbers of (primarily) cortical neurons, is that they are highly rhythmic. While the physiological origins and cognitive function of these rhythms remains unclear, their features are clearly highly labile: spatial location, frequency, and oscillatory power can vary considerably as a function of behavior, cognitive processes, and disease. This suggests that not only the oscillations themselves, but also their fluctuations over time, space, and cognitive state play a key role in brain function. Moreover, multiple frequencies can coexist and interact, fluctuating in a highly correlated manner[1, 2]. Understanding the mechanisms mediating the coexistence of these rhythms, as well as state-dependent changes in their properties, would yield important insight about how collective neural activity and synchronization phenomena, shaped by both sensory and recurrent inputs, mediate neural communication[3]. ‘State’ here simply refers loosely to gross cognitive/perceptual/neural activity regimes, as for example seen in the difference between low-frequency, high-amplitude oscillations observed at rest, and the relatively higher-frequency activity elicited by focused cognitive tasks. In the present paper we opt for the more neutral terms ‘idling’ and ‘active’ (as opposed to ‘rest’ and ‘task’) to indicate these two dynamical regimes. To date only a few models in the literature have sought to explicitly capture transitions between these oscillatory states, and the dependence of certain neural processes on the current state (e.g. [4, 2]).

The majority of neural population models that have been developed to account for the origins of large-scale brain rhythms can be grouped into two broad categories: i) corticalonly and ii) corticothalamic. Cortical-only models typically propose that the oscillatory activity visible in MEG/EEG has its mechanistic origin in interactions between excitatory and inhibitory neurons within a cortical column (e.g. [61, 71, 58]). Corticothalamic models (e.g. [39, 69, 55, 56]) are generally highly similar in overall structure, but differ critically in placing the key excitatory-inhibitory interaction in the thalamus rather than the cortex. These models thus attribute prominent spectral features such as low-frequency oscillations to delayed inhibition in long-range recurrent corticothalamic loops. Given the substantial bodies of empirical data from human and nonhuman physiological recordings supporting each of these two mechanisms, it is highly likely that both play a role in the genesis of large-scale rhythmic activity observed in local field potentials and extracranial electromagnetic fields. Disambiguating the contribution of each to the different features of M/EEG signals, and how they might interact, is a challenging problem, however. Addressing this disconnect is one of the principal aims of the present study.

One of the major points of dispute between cortical-only and corticothalamic model types is the alpha rhythm. Alpha frequency (8-12Hz) oscillations are a hallmark pattern of encephalographic activity[5]. They have been linked to a wide variety of cognitive processes such as perception and attention, and their dynamic features (such as power and frequency) are also closely tied to changes in behaviour[6, 7]. Abnormal alpha activity is also involved in many neurological disorders such as depression, Parkinson’s disease, and Alzheimer’s disease[8, 9, 10]. A broad range of experimental data point to the corticothalamic system as the most likely locus of the dominant alpha-frequency rhythmic activity seen in EEG and MEG[11], as well as the phase relationship between alpha and other faster frequencies. In contrast, gamma frequency oscillations have been robustly tied to intracolumnar excitatory-inhibitory circuit mechanisms and active cortical information processing[12, 13]. It remains an open question, however, how these two types of oscillatory activity (plus associated circuit mechanisms) shape large-scale neural dynamics, functional connectivity, and information integration in a state-dependent fashion.

A key experimental direction for investigating the dynamic properties and functional role of neural oscillations is to study the relationship between endogenous activity and responses to electromagnetic stimulation. This is not only critical for understanding the functional role of brain oscillations in general, but also for improving the efficacy of clinical applications of noninvasive brain stimulation, such as in the treatment of depression[14]. Interestingly, a confluence of experiments with both intra-cranial and non-invasive stimulation have revealed frequency-specific responses, with low-frequency stimulation decreasing the excitability of stimulated tissue[15], and conversely higher frequency stimulation having the opposite effect[16]. Experiments in primates[17] and rodents[18] have indeed demonstrated that thalamic stimulation can be used to either activate or inactivate cortical networks in a frequency-dependent manner, opening new perspectives on the functional manipulation of cortical dynamics by exogenous signals.

To better understand state-dependent changes in oscillatory dynamics, their involvement in inter-area communication, and how they might be controlled by non-invasive stimulation, we present in this paper a novel connectome-based neural mass model that combines cortical and corticothalamic circuit mechanisms in a minimal and parsimonious fashion. In the following sections, we first demonstrate that this model accurately reproduces several key characteristics of measured power spectra and functional connectivity from resting state MEG recordings. We then use the model to study the impact of sensory / neuro-modulatory drive on brain rhythms, and how this serves to switch between low-frequency corticothalamically-driven vs. high-frequency cortically-driven oscillatory regimes. Finally, we show how the model predicts a number of empirical observations in humans and rodents on the relationship between brain state, periodic brain stimulation, rhythmic entrainment of neural activity.

## Results

As detailed in the *Methods*, our full model consists of a network of 68 interconnected nodes, representing brain regions derived from a commonly used parcellation covering most major cortical structures in the human brain. The dynamics of each node is described by a novel extension of the classic Wilson-Cowan (WC) equations[86], which we refer to as the ‘Cortico-Thalamic Wilson-Cowan’ (CTWC) model. Our primary goal was to investigate how state-dependent inputs mediate changes in brain oscillations within multiple frequency bands, and how these spectral fluctuations shape functional connectivity. To do this, we first considered the behaviour of a single isolated network node corresponding to a individual corticothalamic motif. We then moved on to examining collective dynamics and interactions within the whole-brain network.

### Alpha rhythms emerge from delayed recurrent cortico-thalamocortical loops

In examining the dynamics of our corticothalamic model, we first considered the idling state, which we defined as being a state of minimal thalamic drive (see *Methods*) and thus reflecting dynamics in the steady state. Consistent with previous work[4], this system produces a robust alpha rhythm with a spectral peak at approximately 10Hz. In this idling regime, the higher frequency peaks in the power spectrum at beta and gamma frequencies reflect harmonics of the fundamental frequency (alpha), and the background trend in the power spectrum follows a roughly 1/f trend, in line with previous reports[39]. As shown in Figure 1, this model gives a good fit to empirically measured, regionally-averaged MEG power spectra, with all subjects tested showing *R*^2^>=0.6 or higher, and only minor variations in fitted parameter values. Interestingly, we see in empirical MEG data that there are larger differences in power spectra between subjects than between regions within a given subject (data not shown). This observation supports the modelling strategy of choosing a single set of parameters for each subject, and using those for all regions in the network; as opposed to using regionally varying parameter values. We return to the question of spatially varying spectral power below.

**Figure 1:**
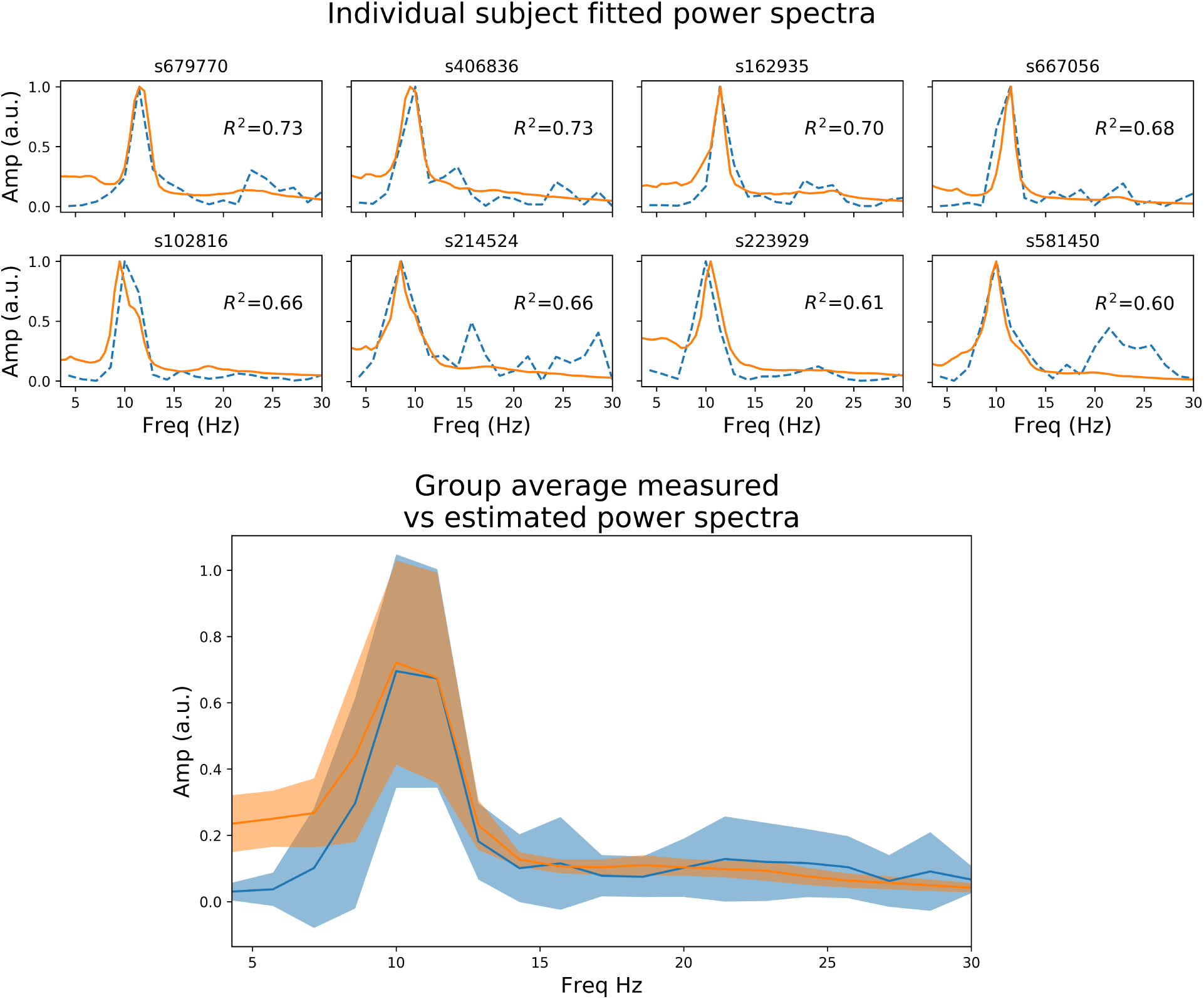
Resting state power spectrum fit to MEG data. *Upper panel:*: Sensor-averaged power spectrum from eight example HCP subjects’ resting state MEG data (orange line), and corresponding simulated power spectrum from the CTWC model (dotted blue line). The simulated activity shows excellent fit to the empirical power spectrum (*R*^2^ between 0.6 and 0.8 in these examples), and accurately captures the alpha rhythm peak frequency in each subject. *Lower panel:* Mean +/-1 standard deviation of the empirical and fitted power spectra for all 10 HCP subjects.

### Phase transition from low-frequency idling to high-frequency active state

Having characterized the dynamics within the idling state and the prevalence of alpha activity, we next asked how increasing the drive to the thalamic populations (either in one or multiple nodes) would impact the spectral properties of cortical activity. To emulate a task or ’active’ state, we thus increased the drive to the thalamic populations (see *Methods*) and observed the resulting behaviour.

We first studied this systematically for a single isolated node. Figure 2 shows trajectories in the 3-dimensional phase space defined by the state variables *u_e_*, *u_i_*, and *u_s_* (representing activity of excitatory cortical, inhibitory cortical, and thalamic specific relay nuclei, respectively), along with time series and power spectra for *u_e_*, which we take as a proxy for M/EEG source activity[39, 58]. The top left panel of Figure 2 shows, the system in the idling alpha-dominated regime, which (consistent with Figure 1) is characterized by a clean and highly stereotyped 10Hz limit cycle. The bottom and top right panels of Figure 2 then show how the system’s dynamics and phase space are modified upon raising the static sensory/neuromodulatory input or drive parameter *I_o_*. We first observe (Figure 2, bottom row) within increasing *I_o_* a gradual destabilization of the resting alpha rhythm, and a transfer of oscillatory power from alpha to higher frequencies. This destabilization is characterized in the 3-dimensional phase space by an increase in the number and regularity of short, rapid excursions (‘twists’) within the alpha limit cycle, which in the time series plots appear as nested higher-frequency ‘ripples’ within the 10Hz base oscillation. Eventually, after a bifurcation point around *I_o_*=1.3 is crossed, the system shifts completely to a noisier, low(er) amplitude gamma-frequency limit cycle, with a clear peak in the power spectrum observed at 30Hz. In line with a confluence of empirical studies[40], this high-frequency component of the power spectrum reflects the fast-paced interplay between excitatory and inhibitory neural populations, and is generated locally within the cortical compartments of each network node. Due to the nature of the corticothalamic circuit motif we considered here, this increased thalamic drive also represents an increased engagement of cortical excitatory and inhibitory populations, that are now recruited for active processing of afferent inputs.

**Figure 2:**
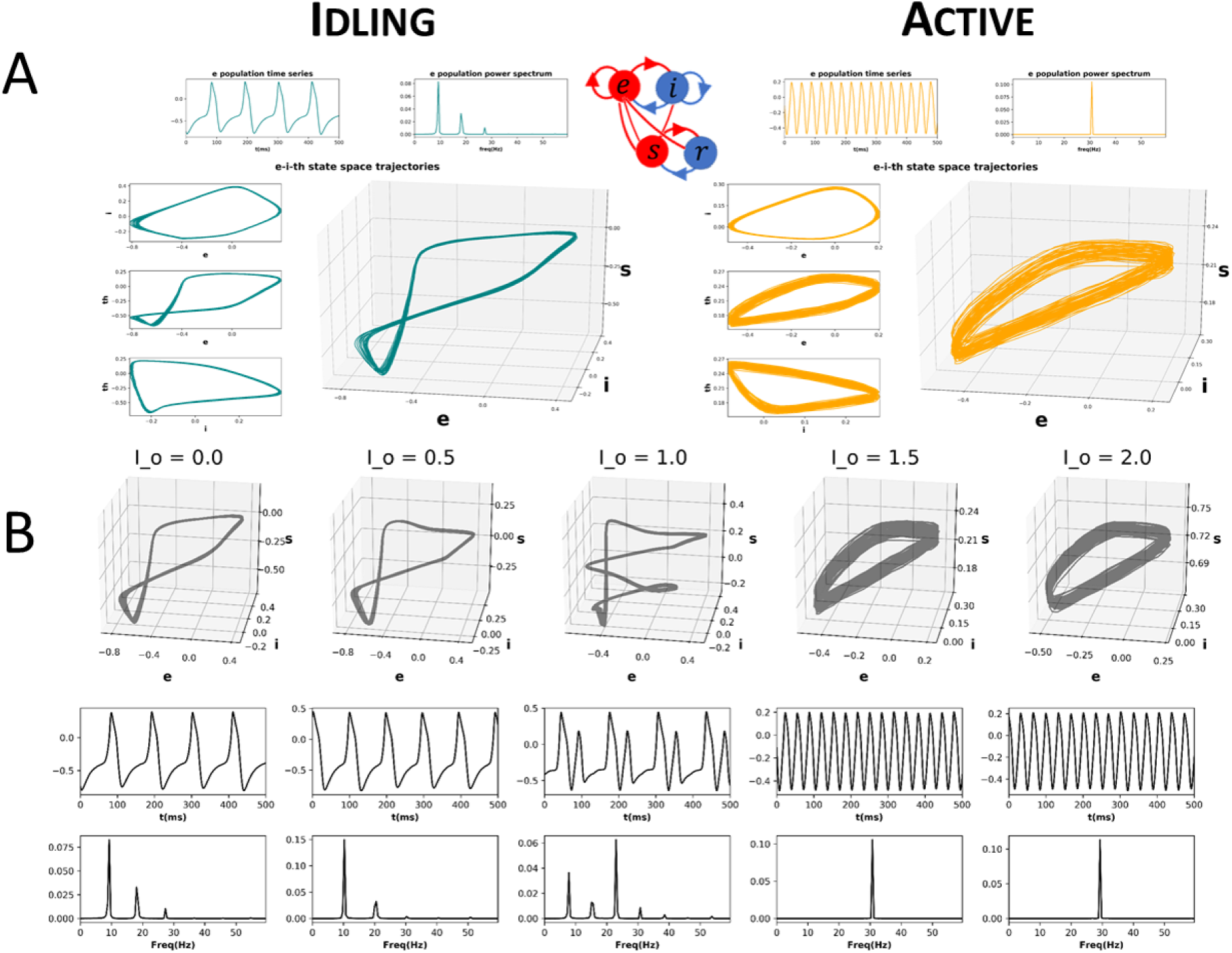
CTWC model phase space trajectories. A) Exemplary phase space trajectories for a single corticothalamic unit in the idling (left; teal) and active (right; orange) regimes. Central 3D plot in each panel shows trajectories in the 3-dimensional phase space defined by the cortical excitatory (***e***), cortical inhibitory (***i***), and thalamic specific relay (***s***) population state variables. Orthogonal 2-dimensional views for each pair of state variables are shown on the left hand side. Panels above the trajectory figures show corresponding time series and power spectra for the ***e*** variable. The idling state regime (*I_o_*=0) is characterized by slow, nonlinear alpha-frequency (8-12Hz) oscillations. Increasing the static sensory/neuromodulatory thalamic drive (here by setting *I_o_*=1.5) induces a phase transition into the active regime, where neural population activity is dominated by gamma-frequency (approximately 30Hz) limit cycle dynamics. B) Progression from idling to active regime. Sub panels show 3D phase plane trajectories, time series, and power spectra for incremental values of *I_o_* between the idling and active states shown in panel A. As the system approaches the bifurcation point (*I_o_* 1.4), the gamma attractor begins to manifest as a ‘twist’ in the alpha limit cycle, which appears in the time series plot as embedded high-frequency ripples on the peak/trough of the oscillation. As *I_o_* continues to be increase, eventually the low-frequency rhythm loses stability and the dynamics switches to a pure gamma oscillation.

### Influence of regionally focal sensory / neuromodulatory drive

We now extend the observations and insights obtained from the single-node case considered in the previous section to the case of whole-brain network behaviour. Figure 3 shows time series, power spectra, and brain-wide plots of the change (Δ) in alpha and gamma power for simulations where *I_o_* is modulated focally for a single node (left V1) in the 68-node network. The suppression of alpha power and enhancement of gamma power with increasing drive is clearly evident in the surface plots and lower power spectrum figure in panel A.

**Figure 3:**
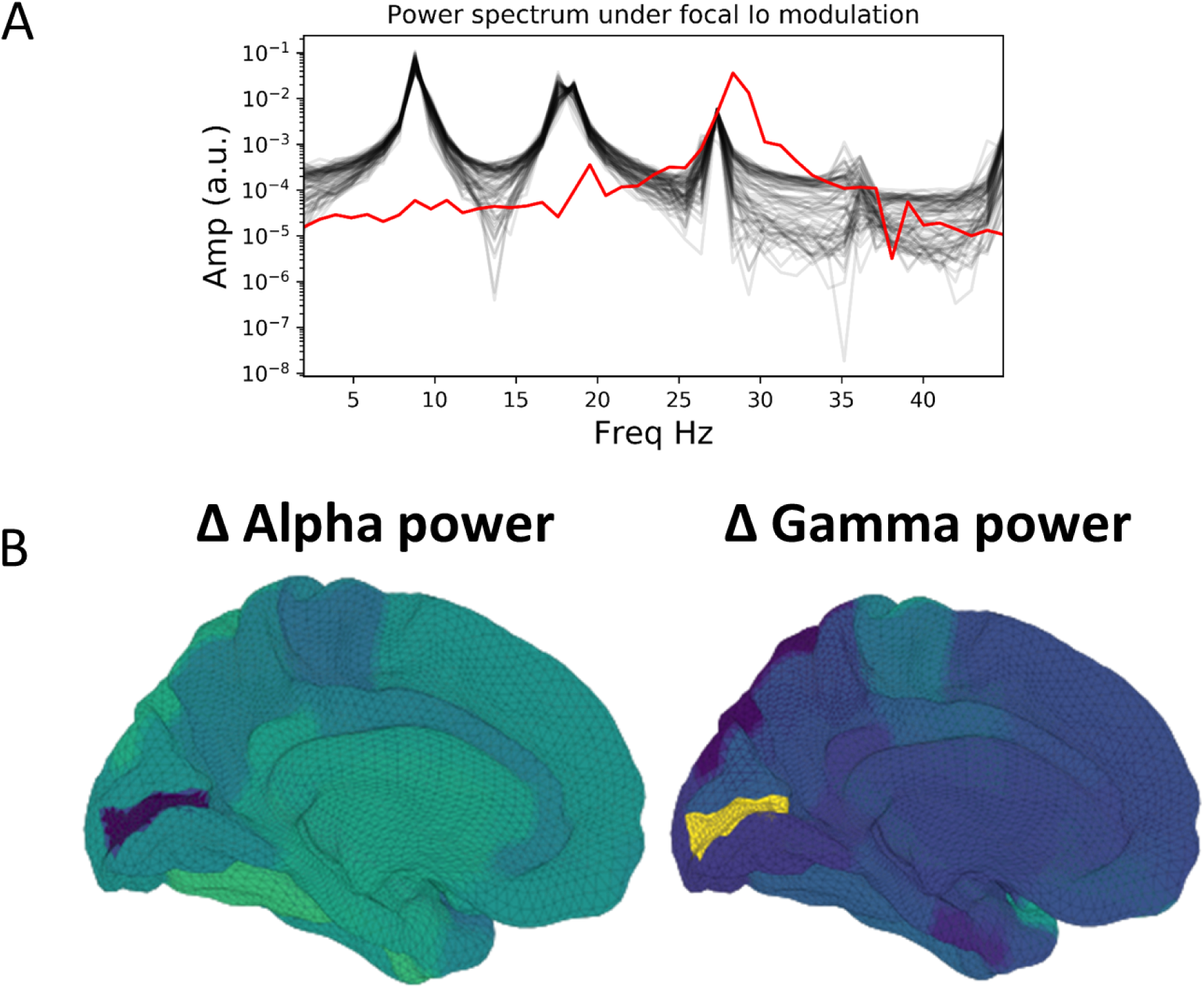
Influence of focal sensory/neuromodulatory drive in a whole brain network. A) Power spectra for baseline values of the tonic thalamic relay nucleus driving term (*I_o_*=0), and for focal increase (*I_o_*=1.5) in left visual cortex (lV1). Red lines show power spectra for the lV1 node; black lines for the other 67 nodes. Note the prominent increase in relative gamma power and decrease in relative alpha power in lV1 when that node’s *I_o_* value is increased. B) Surface renderings of the regional change (Δ) in alpha and gamma power from baseline to active state for all brain regions. Increased sensory/neuromodulatory drive in visual cortex results in suppression of alpha and enhancement of gamma band activity, reminiscent of the patterns routinely observed in M/EEG studies of visual-evoked gamma

### Functional Connectivity

Given the salient differences in oscillatory dynamics observed in the idling and active states, we investigated how these different oscillatory regimes shaped inter-area interactions in a whole-brain network context. To do this, we compared functional connectivity, as measured by amplitude-envelope correlations (AECs) of band-limited power time series, in model-generated time series and empirically measured MEG data.

Heuristically, moving from an isolated node to a network of coupled nodes results in two important changes in the ‘environment’ experienced by each node. First, the overall or time-averaged activity level of a given brain region will be higher when there are inputs from other regions than when there are no inputs. Second, depending on the behaviour of the incoming signals from other regions, that node may experience periodic or otherwise temporally structured driving inputs. This, in turn, may lead to the emergence of synchronization and collective behaviour throughout the system due to processes of entrainment or resonance, possibly also accompanied by bifurcations. As shown in Figure 4, we found idling and active states in the model to be characterized by quite different functional connectivity profiles. The idling state exhibits relatively weaker and spatially non-specific AEC patterns at both alpha and gamma frequencies. In contrast, as the increased static drive *I_o_* pushes the system into the gamma-dominated active state, both alpha- and gamma-frequency AEC matrices increasingly come to display the kind of spatial structure characteristic of empirically measured AEC (as well as by various other M/EEG, fMRI functional connectivity, and indeed anatomical connectivity metrics). Specifically, the active state shows a stronger tendency for spatially nearby regions to show high correlations (as indexed in the AEC matrices by a the ’halo’ of high connectivity values around the leading diagonal), and the classic two-block hemispheric structure with stronger intra-than inter-hemispheric correlations. Interestingly, although the two characteristic frequency regimes within the model are in the alpha- and gamma-ranges, it also captures some properties of AEC outside of these ranges. Figure 5 shows empirical vs. simulated AEC for the full range of classic M/EEG frequencies: delta (0.5-4Hz), theta (4-8Hz), alpha (8-12 Hz), beta (13-30 Hz), and gamma (30-60Hz). As can be seen, moving from low to high frequencies within the active regime is also accompanied by sparser and more spatially structured correlation patterns. It is important to note here that although our model does well at reproducing both resting-state power spectra (Figure and MEG functional connectivity (Figures 4 and 5), the domains in which this success is seen does not entirely overlap. For the power spectrum alone, best fits are achieved at or near ’fully idling’ parameter regime with *I_o_* = 0. For AECs, however, best correspondence with MEG data is achieved in the active regime, with *I_o_* closer to 1.5. We return to this point in the *Discussion*.

**Figure 4:**
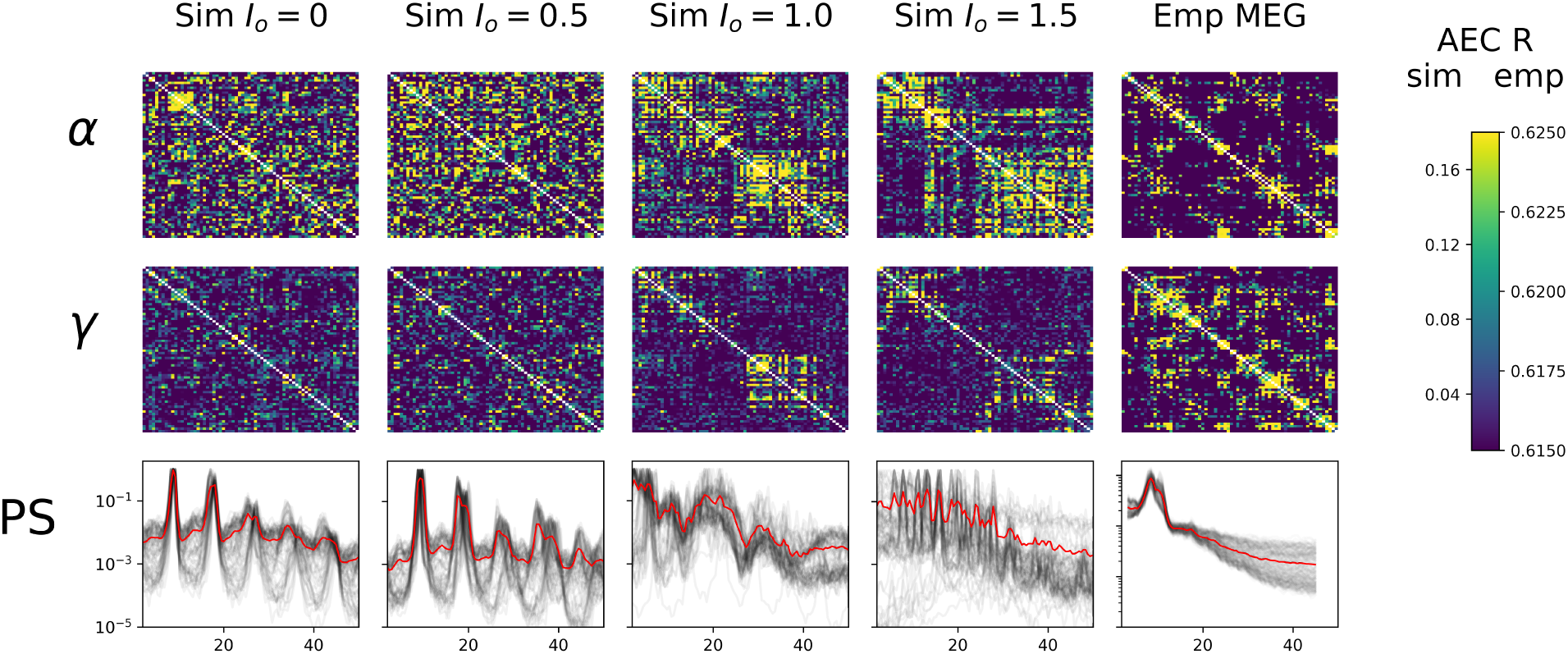
**AEC FC vs.** *I_o_*. *Upper panel* : Gamma-frequency AEC matrices for 4 values of *I_o_* (*I_o_*=0./0.5/1.0/1.5), alongside the empirically-measured MEG gamma-frequency AEC matrix. *Lower panel* : Corresponding power spectra of whole-brain simulated data for these three simulation regimes, as well as for empirically measured MEG data (far right). Black lines show spectra for individual brain regions, thick red line is mean over all brain regions. The simulated power spectra transition from being alpha-dominated at *I_o_*=0 to a noisier and higher-frequency regime at around *I_o_*=1.5.

**Figure 5:**
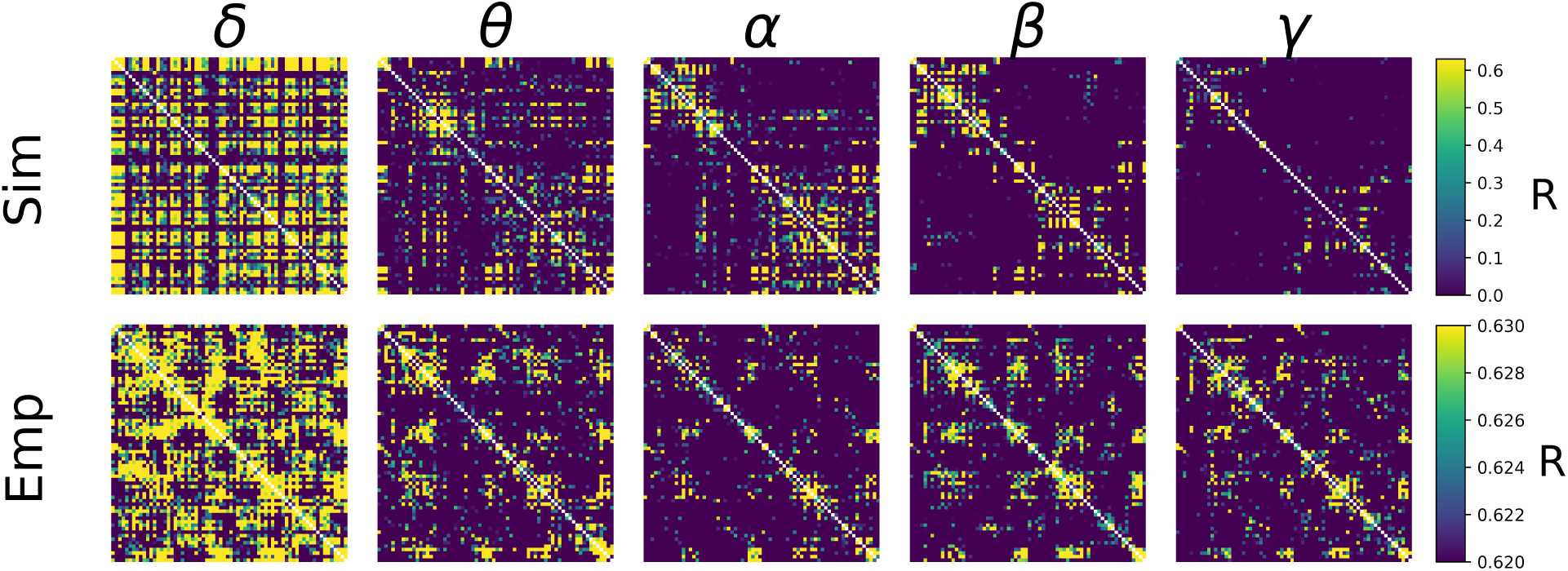
AEC FC vs. Frequency. Shown are AEC FC matrices at five different canonical frequency bands - *δ*(0.5-4Hz), *θ*(4-8Hz), *α*(8-12Hz), *β*(13-30Hz), and *γ*(30-45Hz) - from empirical MEG data (top row), and from simulations (bottom row). In both simulated and empirical data, lower frequencies (*δ* and *θ*) show less spatial specificity and more tendency towards random connectivity patterns. Note that the more compressed AEC range in empirical than simulated AEC data is due to the application of orthogonal leakage correction[89] in analyses of MEG data.

Our findings described thus far have shown that active and idling states are characterized by different spectral signatures, and that functional connectivity is differentially expressed in a frequency-specific way in these two states. Next, we examined the effects of periodic stimulation on ongoing cortical activity. That is, we asked: can the temporal structure neural activity be tuned by exogenous signals in a frequency-specific way?

### Susceptibility to entrainment by exogenous stimulation is state-dependent

Having characterized idling and active states, their dominant spectral features and how they impact functional connectivity, we investigated how *exogenous* periodic stimulation shapes the power spectrum of the system and engages ongoing oscillations. Numerous studies over the last few decades have used stimulation paradigms of various kinds to access circuit function and interfere with neural communication[42, 43, 44]. One of the most robust findings is that entrainment of ongoing brain oscillation is state-dependent, and that susceptibility to control is tuned by ongoing brain fluctuations - an effect that has also been reproduced with modelling[45, 46] and shown to involve stochastic resonance[47]. Given the ability of our model to switch between different states and express multiple frequencies, we subjected cortical populations to exogenous periodic stimulation and monitored the spectral response. Specifically, we again studied an isolated cortico-thalamo-cortical motif (i.e. a single network node), and computed the peak power and frequency as a function of stimulation intensity and frequency. Through this process, we identified resonances and entrainment regimes (so-called *Arnold Tongues*) and thus measured the susceptibility of our model to entrainment. While oftentimes confused with one another, *resonance* refers to the enhancement of power when the stimulation frequency is in the vicinity of the system’s natural frequency, while *entrainment*, refers to the phase locking of the system’s response to the driving frequency[47]. As shown in Figure 6, idling and active states exhibited significant differences in their responses to stimulation and susceptibility to entrainment. Narrower Arnold Tongues were observed in the idling state compared to the active state, indicating that the suppression of alpha power in the active state facilitates phase locking of intrinsic dynamics with the stimulation signal. Specifically, only high intensity stimulation would provoke a shift in the peak frequency in the idling state. In the active state, the prominent gamma oscillations were easily suppressed and replaced by the frequency of the driving stimulus. This is in line with converging evidence indicating that intrinsic attractors limit the effect of perturbations, while irregular or high frequency content is more malleable[4].

**Figure 6:**
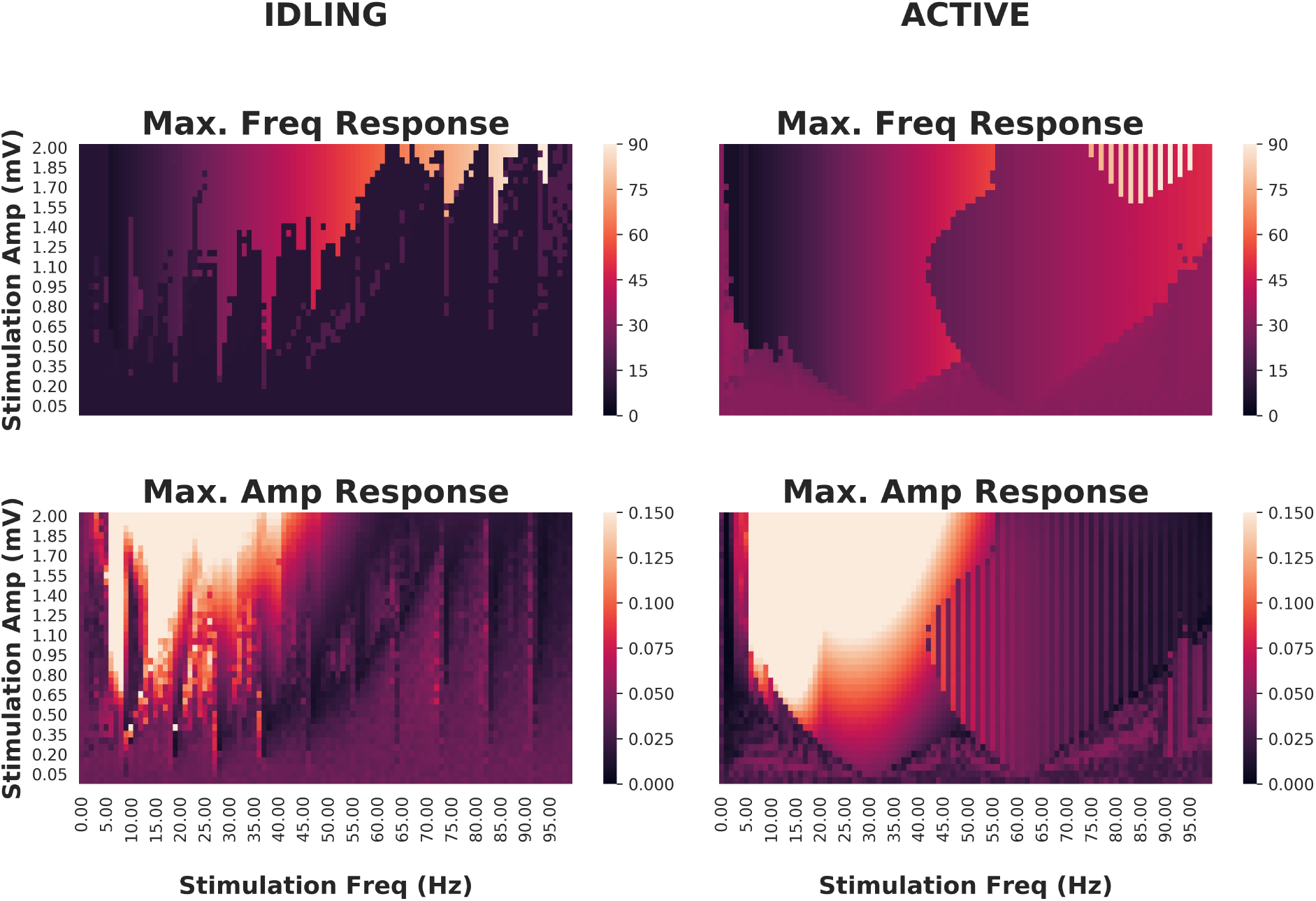
Effects of periodic brain stimulation on corticothalamic loop dynamics. *Top row:* Maximum frequencies displayed by the cortical excitatory population of an isolated cortico-thalamocortical loop (CTWC model, single node) in response to periodic (sine wave) stimulation of varying amplitudes (y axes) and frequencies (x axes). In the idling regime, an Arnold Tongue structure is clearly seen centred on the natural frequency (approximately 10 Hz): As the stimulation frequency moves away from the natural frequency, greater stimulation amplitude is required to achieve entrainment at the stimulation frequency. In the active regime, a broader and shallower Arnold Tongue structure is again seen, centred on the natural frequency (this time approximately 30Hz). Compared to the idling state, entrainment at the stimulation frequency is easier to achieve (requires lower amplitude stimulus) in the active than the idling regime. *Bottom row:* Maximum amplitudes displayed by cortical excitatory populations. Here again the amplitude response patterns match quite closely the Arnold Tongues seen in the maximum frequency responses.

## Discussion

The aim of the present study was to investigate the mechanisms underlying state-dependent changes in oscillatory activity at the whole-brain scale, as well as the influence of fluctuations in spectral activity on functional connectivity. We have presented a novel connectome-based neural mass model that combines the two primary rhythmogenic mechanisms typically studied in large-scale brain network modelling: intracolumnar microcircuits and corticothalamic loops. This is an extension of previous work, that studied the behaviour of the basic corticothalamic motif in isolation[48]. Here we have embedded this corticothalamic unit into a whole-brain network, with anatomical connectivity derived from diffusion MRI tractography.

Our model reproduces a variety of known features of human M/EEG recordings, including a 1/f spectral profile, spectral peaks at canonical frequencies, and functional connectivity structure that is shaped by the underlying anatomical connectivity. Using this model, we have studied how thalamic drive mediates a shift in oscillatory regime, provoking a transition between alpha and gamma dominance in the power spectrum, and found that these oscillations have a differential impact on functional connectivity patterns. We found that spatially structured inter-area functional connectivity (as measured by band-limited power amplitude envelope correlations), particularly at higher frequencies (gamma, beta, and alpha to a lesser extent), are a hallmark of the active state. To better understand how these state- and frequency-specific dynamics are impacted by exogenous stimulation, we applied cortical periodic stimulation of various amplitudes and frequencies, eliciting endogenous resonances both across the corticothalamic loop and within cortex. Our analysis confirms that, as compared to the idling state, the active state is more susceptible to entrainment by exogenous signals, as it shows wider and shallower Arnold Tongues. In contrast, the idling state’s deep and narrow Arnold Tongues indicate that the system has a strong preference for its natural frequency when in this regime, and will respond only to exogenous signals close to that frequency or its harmonics.

### Relation to previous work

The work presented here builds on previous work of several authors in a number of ways. Most directly, the isolated CTWC neural mass model (without the whole-brain white matter connectivity introduced here) was recently introduced in [48]. Previous to that we have also studied resonance behaviour, response to stimulation, and state-dependence in corticothalamic circuits and generic feedback oscillators[45, 65, 64]. We emphasize however that the core mathematical and conceptual component of the CTWC model presented in the present paper and in our earlier work - namely the generation of slow M/EEG rhythms through a delayed inhibitory cortico-thalamo-cortical recurrent circuit, has been used extensively by multiple groups for several decades. One of the largest and most comprehensive bodies of work on this is due to P. Robinson and colleagues, beginning with the introduction in [79] of a PDE wave equation reformulation of the integro-differential cortical neural field model of [83], drawing on earlier work of [11], [66], and others. This model was then augmented with thalamic reticular and relay nuclei and their recurrent connections with the cortex[39], and the resultant corticothalamic neural field model has been studied extensively over the past two decades - both analytically and numerically, and in partial differential, ordinary differential, and lin- earized equation forms, as well as being extended into the domains of epilepsy, Parkinson’s, sleep and arousal, plasticity, and brain stimulation (e.g. [39, 69, 67, 70, 84, 85, 82, 81, 68]). Our approach in the present paper differs from this family of models in two key ways. First, rather than the second-order equations of motion for the time-evolution of membrane voltage used by Robinson and many others[11, 61, 71], we began with the classic Wilson-Cowan equations[86] to describe local interactions between excitatory and inhibitory neural populations in a cortical region. Second, rather than the using an integro- or partial-differential equation formulation of a continuum neural field to represent spatio-temporal propagation of activity across the cortex[79, 66, 51, 74, 72, 73], here we chose to follow the connectome-based neural mass modelling methodology[75, 76, 77, 26, 63, 20, 54] of defining a discrete network of point-process neural masses, interconnected via long-range white matter fibres whose density was estimated from non-invasive diffusion MRI tractography. This combination of the cortico-thalamocortical circuit with the large-scale anatomical connectivity bears some similarity to the work of some other authors (e.g. [25, 56, 55, 80]), but the present study is the first to apply this directly to the key questions of state-dependence, alpha suppression, functional connectivity, stimulation, and their relation to empirical M/EEG data. Notably, this network-based approach allowed us to harmonize the analysis of functional connectivity in simulated and empirical MEG data. In this we followed the approach of [36] and [87] in our use of the bandpass-filtered amplitude envelope correlations[78, 37], and that line of work is perhaps the closest of recent modelling studies to the present one. In [36], Abey-suriya and colleages studied the role of inhibitory synaptic plasticity in a connectome-based network of Wilson-Cowan equations. As in the present study, these authors evaluated their model in terms of its ability to accurately reproduce empirically measured MEG AEC matrices (although they restricted their focus to only to alpha-frequency AECs). The relatively simpler (as compared with our new CTWC) model used by these authors consisted of a cortical Wilson-Cowan ensemble, tuned to have a natural frequency in the alpha range. This stands somewhat in contrast to our new model, which features a *gamma* frequency-tuned Wilson-Cowan ensemble, combined with an *alpha* frequency-tuned cortico-thalamocortical motif. This additional two-component structure allows our model to exhibit more complex behaviours, such as alpha-mediated inhibition and state-switching, as well as a rich repertoire of potential oscillation and frequency-specific synchronization patterns. The question of whether and to what extent human M/EEG alpha activity is generated by corticothalamic (as in e.g. the present study and much of the above-cited work by Robinson and colleagues), or within intracortical microcircuits (as in e.g. [36], [58], [71]) remains a live and important one however. Recent years has also seen growing interest in a third potential type of system-level (low-frequency) rhythmogenic mechanism which can be broadly described as *network eigenmodes*[51, 79, 72, 63, 62]. The proper evaluation and assessment of these hypotheses around cortical rhythmogenesis shall most likely require a close interaction between novel empirical work and hypothesis-generating computational models to properly settle. It is also important to bear in mind here that there is no a priori reason (apart from explanatory parsimony) to suppose a single mechanism for generation of rhythms[51]. Indeed, it may be functionally advantageous for the brain to generate the same frequency through a variety of mechanisms. If this were determined to be the case, then interaction across different frequency-generating mechanisms would be a key question for future work.

### The alpha rhythm as a suppression mechanism

The transition from idling to active state in our model is initiated by the gradual increase of a tonic sensory/neuromodulatory drive term, *I_o_*, that effectively hyperpolarizes the thalamic relay nucleus, and thereby destroys the slow 10Hz alpha rhythm generated by the cortico-thalamocortical loop. Once the alpha oscillation is removed in this way, the gamma rhythm generated by intracortical excitatory-inhibitory interactions comes to the fore. One interpretation of this phenomenon is that alpha resonance, mediated by corticothalamic loops, plays an inhibitory role - through which slow oscillatory corticothalamic activity suppresses and dominates higher frequency cortical activity. This alpha-as-suppression-mechanism theory speaks to a major question in the field of M/EEG cognitive neuroscience: what is the functional role of alpha? Specifically, the enhancement of alpha activity during disengagement of the cortical network (such as during quiescence, sleep, anaesthesia, and withdrawal of sensory stimulation) suggests that alpha oscillations implement a functionally inhibitory signal, and represent a top-down shift towards internal encoding through suppressing the activity of task-irrelevant areas[49]. In contrast, faster frequencies, such as those found in the beta and gamma range, are found in states of arousal and sensory recruitment, suggesting a positive, excitatory role of faster neural oscillatory states. In our model, the less spatially-resolved structure of functional connectivity in the alpha vs. the gamma range - at all *I_o_* values, but particularly for *I_o_*>=1.4 - does support this perspective. From this point of view, a key feature of our model is its characterization of the relationship between corticothalamically-generated and cortically-generated rhythms. In particular, the corticothalamic alpha dominates in the idling state, and can be understood as suppressing the intrinsic rhythmic activity in the cortical ensemble, which can be ‘released’ with sufficient sensory or neuromodulatory drive. This simple circuit mechanism therefore captures a widely used theoretical concept in M/EEG cognitive neuroscience concerning the functional role of alpha activity. On this account, alpha acts as a mechanism for selectively gating and attentionally biasing sensory inputs. This phenomenon is also observed in EEG studies on the effects of anesthesia, where low frequency activity becomes increasingly dominant with higher doses of propofol[50]. This effect is observed concurrently with apparent attenuation of sensory inputs, for example in reduced amplitude and increased latency of somatosensory evoked potentials (SEPs). Recent work in mouse models has also shown that driving thalamic circuits with alpha-frequency activity causes widespread depression of cortical activity; whereas stimulating at higher frequencies (e.g. gamma) causes widespread increase in both baseline activity and the spatial spread of the stimulation influence[18].

Interestingly, in our analyses we observed that the active-state model AEC patterns actually showed closer resemblance to empirical resting-state MEG AEC patterns than the idling-state AEC patterns. This is somewhat unexpected because resting-state MEG power spectrum was unequivocally better fit by a CTWC model in the idling, alpha-dominated regime. This result suggests that in the brain, during the rest or idling state, alpha power is strong and AEC functional connectivity is largely random. In contrast, in the active state, alpha power is relatively weaker, and AECs are more local and segregated. Functional connectivity is thus facilitated in the high-drive state, when the alpha-generating loop is inhibited, and dynamics are driven by cortico-cortical E-I interactions. In the state of low-drive, the alpha rhythm is highly prominent and functional connectivity is largely asynchronous. In the state of high drive, the alpha rhythm has been suppressed, and functional connectivity is high. Together, these observations suggest that the alpha rhythm plays a suppressing role in large-scale brain dynamics. We hypothesize that this may be a general feature of alpha activity - this indicates that regional communication is facilitated by being in the active state, and that there perhaps a constant interplay and balance between the idling state and the active state.

### Conclusions and future directions

To conclude: we have developed a novel whole-brain connectome-based neural mass model that incorporates corticothalamic and intracortical rhythmogenic mechanisms. This model reproduces qualitatively multiple features of MEG-measured neural activity. Importantly, our model also lends some insight into the way that cortico-thalamically-generated alpha rhythms could play a functional role in the organization of brain dynamics, by suppressing high-frequency cortical activity associated with cognitive engagement and information processing. Future work shall investigate further questions of subcortical parcellation and integration, model fitting, and compare alternative rhythmogenic mechanisms directly against each other. Importantly, future work should also investigate the significance of intersubject variability in anatomical connectivity on network dynamics. Although we demonstrated here our model’s ability to fit individual subjects’ power spectra through small variations in thalamic kinetic parameters, it was beyond the scope of the present study to incorporate individualized anatomical connectivities. One of the exciting and promising aspects of connectome-based neural mass modelling is the possibility of constructing individualized computational models using a subjects’ own diffusion MRI tractography. However at this point in time the extent to which this does actually deliver improvement in computational model accuracy remains an open question for the field (for recent work relevant to this, see [36, 60]). Finally, we emphasize that neither our specific CTWC model, nor the broader alpha-as-suppression-mechanism concept, constitute a universal account of all alpha-frequency rhythms seen in the M/EEG or other recording modalities. Indeed we consider the most likely scenario to be that multiple, dissociable mechanisms contribute independently a proportion of the information and measured signal in that part of the frequency spectrum[51]. Here we have, building on previous work, made we believe some progress in characterizing the dynamic properties of one of these candidate mechanisms.

## Methods

Our modelling approach follows the now-standard whole-brain connectome-based neural mass modelling paradigm[75, 77, 19, 20], where dynamic units are placed at node locations as defined by a grey matter parcellation, and coupled with an adjacency matrix (anatomical connectome) defining the presence and associated strengths of long-range white matter fibres interconnecting region pairs. The anatomical connectome used in the present study, derived from group-average tractography streamline counts, was constructed from analyses of the human connectome project (HCP) WU-Minn consortium diffusion-weighted MRI (DWI) corpus[21, 22]. For details of this, see the below section *DWI data analyses*.

In the model, activity at each node is driven by background noise and/or exogeneous stimulation. Complete mathematical formulation and implementation details are given in the section *Corticothalamic model.* Simulated nodal time series from the model can be understood as approximations of regionally averaged source-space MEG signals. To assess the performance of the model in reproducing key features of empirically measured human brain dynamics, we additionally conducted new analyses of the HCP WU-Minn resting-state MEG corpus[23]. These are described in the *MEG data analyses* section.

### Corticothalamic model

Following other authors [39, 24, 25], we employ a model for neuronal dynamics at each node that incorporates both cortical and thalamic neural populations. The model describes a four-component cortico-thalamo-cortical motif, consisting of excitatory (**u***_e_*) and inhibitory (**u***_i_*) cortical neuronal populations, coupled to thalamic reticular (**u***_r_*) and specific relay (**u***_s_*) nuclei (Fig. 7). Both relay and reticular nuclei receive inputs from the cortical excitatory population, following a corticothalamic conduction delay *τ_ct_*. However only the relay nucleus sends excitatory input back to the cortex; again received following a delay *τ_ct_*=*τ_tc_*. The reticular nucleus, which is widely known to have an inhibitory influence of other thalamic regions[59], plays a similar role to the cortical inhibitory population, inhibiting the relay nucleus and thereby generating oscillatory dynamics.

**Figure 7:**
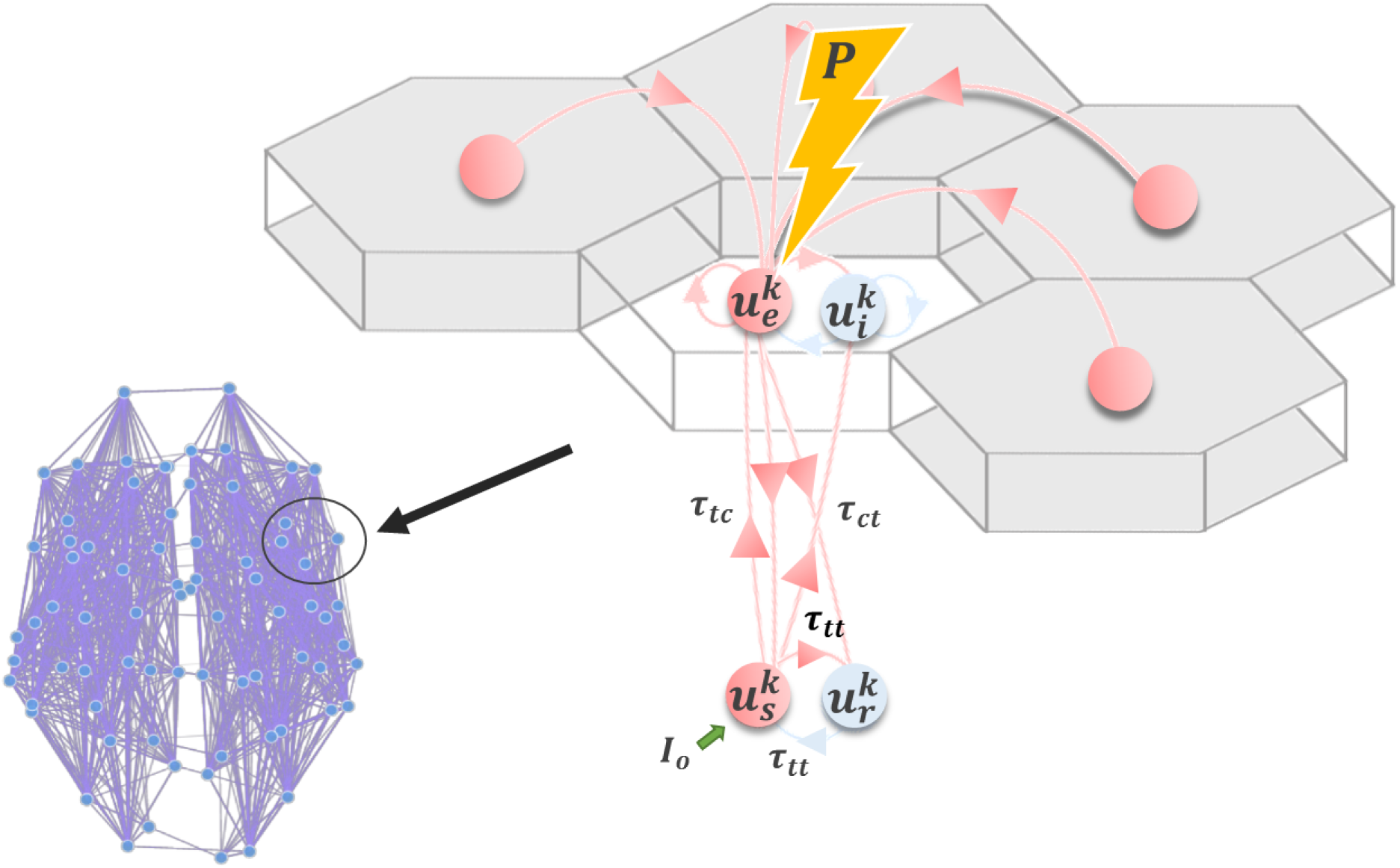
Corticothalamic model. Schematic of the corticothalamic model structure. Cortical (*u_e_*, *u_i_*) and thalamic (*u_s_*, *u_r_*) populations interact through a delayed feedback loop. Entrainment of the network activity through electromagnetic stimulation *P* applied to *u_e_* depends on the amplitude and frequency of the stimulation pulse, as well as the network state, controlled by *I_o_*.

As defined, our node-level model consists of a Wilson-Cowan oscillatory neural population, embedded in a delayed inhibitory feedback loop mediated by corticothalamic and thalamocortical connections. The full network-level model thus consists of a set of *N* such local units of this kind, coupled using the connectivity matrix **W** (anatomical connectome). The system of stochastic delay-differential equations governing the time-evolution of neural activity within the network can be summarized as follows:

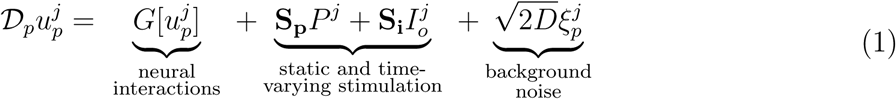

where the temporal differential operator 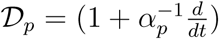 incorporates population time constants *α_p_*, and 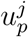 refers to the mean somatic membrane activity of the neural population *p ∈ {e, i, r, s}* within one cortico-thalamic module *j* across the brain-scale network of *N* =68 nodes. Irregular and independent fluctuations are also present in the network, modelled by the zero-mean Gaussian white noise 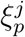 processes with standard deviation *D*. The neural interaction term 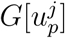 in Eq. 1 can be further broken down into

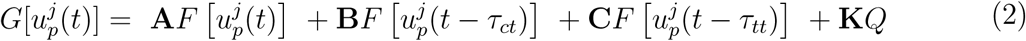

where the matrices

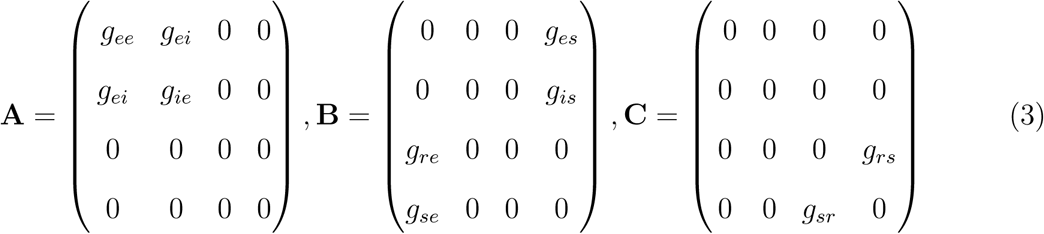

respectively specify the gains (connection strengths) of intracortical, corticothalamic and intrathalamic interactions within a node. Intrathalamic and corticothalamic/thalamocortical connections are retarded by conduction delays *τ_ct_*=20ms and *τ_tt_*=5ms, respectively. The matrix

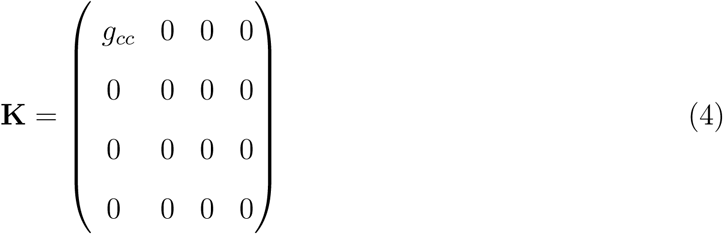

specifies the global gain applied to all afferent activity *Q* arriving from other cortical neural populations. In the present model we assume for simplicity that afferent activity only impacts on the cortical excitatory population *u_e_*; and so only the upper left entry in **K** is nonzero. The afferent activity in *Q* is a time-delayed summation of *u_e_* at all other nodes in the network

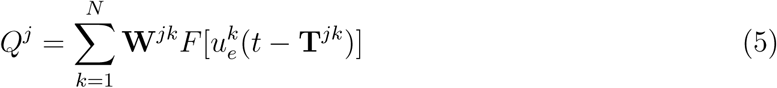

where **W** and **T** are cortical white matter connectivity and conduction delay matrices, both of which are derived from empirical diffusion-MRI tractography reconstructions (see below). For the latter, the cortico-cortical conduction delay matrix **T** = **L***/cv* is calculated from a matrix of measured (average) fibre tract lengths **L**, assuming a fixed conduction velocity *cv*=4m/s. The sigmoidal response function *F* in Eqs. 2 and 5 specifies the nonlinear response of a neural population to incoming inputs as follows

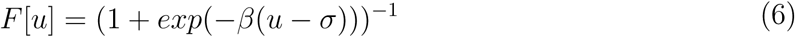

The matrices

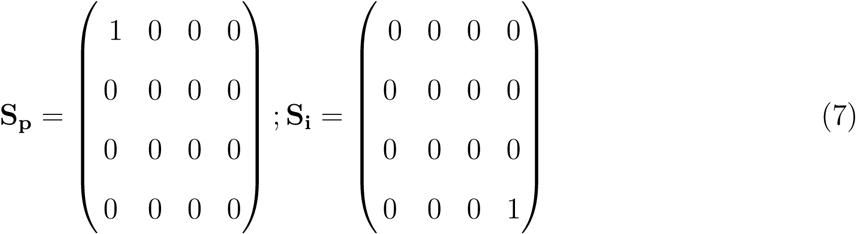

in Eq. 1 parametrize the impact on the four subpopulations *e, i, r, s* within a node of the time-varying exogenous input *P* (representing periodic brain stimulation such as rTMS or TACS) and static input *I_o_* (representing here state-dependent sensory/neuromodulatory drive). Again, in the present study we only consider exogeneous inputs to impact the cortical excitatory populations, and so only the upper left entry in **S_p_** is nonzero. Similarly, *I_o_* is for present purposes only considered to impact the thalamic relay nucleus, and so only the lower right entry of *S_i_* is nonzero. The exogeneous periodic signal *P* here is given by the simple sinusoidal function

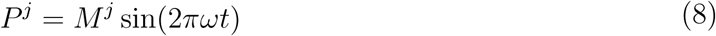

with frequency *ω* and intensity *M*. The constant state-dependent drive 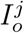 to thalamic relay populations serves as a control parameter indexing idling vs active states (see below). This static input current can be thought of as a tonic level of sensory (e.g. visual) drive, although it could also reflect a static influence of ascending (e.g. noradrenergic) neuromodulatory drive, reflecting the level of engagement in a perceptual or cognitive task. Irrespective of its cause, the idling or rest-like state is defined as the dynamics resulting from setting 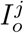=0; i.e. in the absence of this constant thalamic input. The active state, in contrast, is defined by a greater engagement of thalamic nodes, and hence 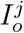>0 for active nodes. In both of these cases, nodes within the network may be differentially recruited by a given task, thus being activated while others remain inactivated. This represents an intermediate point between the extreme cases where all nodes are either active or inactive.

With the described structure, and right choice of parameters, our system generates alpha (8-12Hz) oscillations due to the presence of delayed inhibition, as well as gamma (30-120Hz) oscillations resulting from the cortical activity and interactions, and also in a limited domain of parameter space shows coexistence of both of these features. As has been demonstrated previously [48], increasing the thalamic drive parameter past a critical point triggers suppression of resting state alpha oscillations, and results in a greater susceptibility of cortical neural populations to entrainment by exogenous inputs or noninvasive stimulation. In addition, this transition to the active state is accompanied by an increase in high-frequency (i.e. gamma) activity. As such, the thalamic drive can be seen as a control parameter, controlling the power of alpha and gamma oscillations, as well as tuning the response to exogenous inputs.

Nominal parameter values and definitions from the above-specified system of equations are summarized in Table 1. The system was numerically integrated using a stochastic Euler-Maruyama scheme, implemented in Python. Simulations were carried out on an 8-core Ubuntu 14.04 machine. Run time scaled approximately linearly: each 2-second simulation ran in approximately 2 seconds real time. All code and processed data used in this study is freely available at https://github.com/GriffithsLab/ctwc-model, along with additional notes and comments. A version of the model has also been developed for direct use within The Virtual Brain modelling and neuroinformatics platform (TVB; www.thevirtualbrain.org)[26, 27]). Our model produces regional time series for each network node, as specified by the anatomical parcellation. These represent the collective activity of neural populations within that region, and as such correspond to signals estimated from MEG source reconstruction. Subsequent power spectrum and functional connectivity analyses of simulated activity time series therefore proceeded identically to that for MEG data, and are described in the *MEG data analyses* section below.

**Table 1:** Model parameters.

| Name | Unit | Nominal Value | Description |
| --- | --- | --- | --- |
| $a_e$ | $ms$ | 0.3 | Cortical excitatory population time constant |
| $a_i$ | $ms$ | 0.5 | Cortical inhibitory population time constant |
| $a_s$ | $ms$ | 0.2 | Thalamic relay nucleus time constant |
| $a_r$ | $ms$ | 0.2 | Thalamic reticular nucleus time constant |
| $i_e$ | $mV$ | -0.35 | Cortical excitatory population constant input |
| $i_i$ | $mV$ | -0.3 | Cortical inhibitory population constant input |
| $i_s$ | $mV$ | 0.5 | Thalamic relay nucleus constant input |
| $i_r$ | $mV$ | -0.8 | Thalamic reticular nucleus constant input |
| $\tau_{(ct/tc)}$ | $ms$ | 20 | Corticothalamic / Thalamocortical conduction delay |
| $\tau_{tt}$ | $ms$ | 5 | Thalamo-thalamic conduction delay |
| $I_o$ | $mV$ | 0. | Static sensory/neuromodulatory drive |
| $dt$ | $ms$ | 0.1 | Integration step size |
| $w_{ee}$ | | 0.5 | Excitatory-excitatory gain |
| $w_{ei}$ | | 1 | Excitatory-inhibitory gain |
| $w_{ie}$ | | -2. | Inhibitory-excitatory gain |
| $w_{ii}$ | | -0.5 | Inhibitory-inhibitory gain |
| $w_{er}$ | | 0.6 | Excitatory-reticular gain |
| $w_{es}$ | | 0.6 | Excitatory-relay gain |
| $w_{si}$ | | 0.2 | Relay-inhibitory gain |
| $w_{se}$ | | 1.65 | Relay-excitatory gain |
| $w_{rs}$ | | -2. | Reticular-relay gain |
| $w_{sr}$ | | 2. | Relay-reticular gain |
| $D_{(e,i,r,s)}$ | | 0.0001 | Noise standard deviation for all populations |
| $g$ | | 0.9 | Global connectivity scaling factor |
| $\beta$ | | 20. | Activation function gain parameter |
| $\sigma$ | | 0. | Activation function threshold parameter |

### DWI data analyses

The anatomical connectivity matrices used in this paper were constructed using diffusion- and T1-weighted MRI data from the HCP WU-Minn consortium[28, 21, 22]. For detailed descriptions of the MR acquisition parameters and processing pipeline, see [21, 22]. The HCP WU-Minn corpus consists of multimodal imaging and behavioural data from 1200 healthy, young (ages 20-40) subjects. The tractography analysis described below was applied to a 700-subject subset of the full sample; and the connectivity matrix used for simulations in the present paper was calculated from an average over these 700 subjects.

The HCP WU-Minn minimal diffusion pipeline[21] consists of gradient nonlinearity correction, eddy current correction, boundary-based registration and reorientation of diffusion data to the T1 image, and gradient vector rotation. The outputs of this preprocessing pipeline were the starting point for our diffusion data analyses. Using the minimally preprocessed diffusion data, we performed whole brain deterministic tractography reconstructions using the Dipy software library[29], following a methodology modelled closely on that of [30] and [31]. ODFs were computed at each white matter voxel using a DSI tissue model. Streamlines were initiated from 60 regularly-spaced grid points within each voxel on the grey-white matter interface (as determined from coregistered freesurfer surfaces), and propagated using the EuDX algorithm[32]. Streamlines not terminating at the grey-white matter interface, or having lengths greater than 250mm or less than 10mm, were discarded. Subjects’ streamline sets were segmented using the Lausanne scale-1 parcellation[30, 33], computed individually for every subject from their freesurfer reconstructions using algorithms from the connectome mapping toolkit[33]. All surface-based parcellations were then converted to image volumes and resliced to diffusion space for streamline segmentations. For each parcellation, the interconnecting streamlines for every ROI combination were determined using a logical AND operation. Each segmented streamline set was counted and its average length computed, resulting in streamline count and length matrices for each subject. The simulations described in the present paper were computed using group-average tract length matrices (divided by conduction velocity to convert to conduction delay), and group-average streamline count matrices, with the latter first being log-transformed to adjust for the DWI tractography over-estimation bias[36].

### MEG data analyses

MEG analyses were performed using 10 randomly selected subjects from the HCP WU-Minn corpus[23], using the MNE software library[35, 34]. The specific analyses done were based on a modified version of the analysis pipeline developed by Engemann and colleagues (https://github.com/mne-tools/mne-hcp), which implements a full source space analyses, beginning with the HCP preprocessed sensor-space data. Key outcome variables from this pipeline for the present study were whole-brain functional connectivity matrices and spectral power maps, derived from regional source time series estimates. We opted to implement a complete analysis here rather than use the high-level pipeline outputs provided with the HCP WU-Minn corpus, as we needed complete control over the process. In particular, we needed to a) use the same parcellation in the MEG as in the tractography analyses, and b) ensure identical analyses were done on empirical and simulated MEG regional time series. Regarding the first of these: as in the tractography analyses, the parcellation used for MEG analyses was the Lausanne2008 scale 1 - but with 10 subcortical nodes (brainstem, basal ganglia, thalamus) excluded. Note this is in fact identical to the freesurfer *aparc* parcellation (but reordered and renamed).

Source time series were extracted for all vertices within a parcel using an L2 minimum-norm inverse solution and averaged, yielding one representative time series per parcel. To maximize robustness of these signals, this was operation was repeated five times, with 30-second windows each[23]. Subsequent analysis of these regional time series proceeded identically for both the empirical and simulated MEG data. We first computed power spectra for each region using Welch’s method. We then studied functional connectivity within the system using the band-limited power Pearson correlation (BLPC) method[37, 38]. For this, regional time series from each of the 5 windows were bandpass-filtered into six canonical frequency bands: delta (0.5-4Hz), theta (4-8Hz), alpha (8-12Hz), beta (12-30 Hz), low gamma (30-50 Hz) and high gamma (60-80 Hz)[37]. Pearson correlations between the bandpass-filtered time series were computed, and averaged over the 5 windows. Finally, these BLPC matrices at each frequency band were averaged over subjects. Because our simulations used a normative (rather than subject-specific) anatomical connectivity, these analyses were conducted only once on the simulated MEG data, and this was compared to the group-averaged MEG data to evaluate the performance of the model.

